# Predictability and persistence of prebiotic dietary supplementation in a healthy human cohort

**DOI:** 10.1101/222919

**Authors:** Thomas Gurry, HST Microbiome Consortium, Sean M Gibbons, Le Thanh Tu Nguyen, Sean M Kearney, Ashwin Ananthakrishnan, Xiaofang Jiang, Zain Kassam, Eric J Alm

## Abstract

Dietary interventions to manipulate the human gut microbiome for improved health have received increasing attention. However, their design has been limited by a lack of understanding of the quantitative impact of diet on a host’s microbiota. We present a highly controlled diet perturbation experiment in a healthy, human cohort in which individual micronutrients are spiked in against a standardized background. We identify strong and predictable responses of specific microbes across participants consuming prebiotic spike-ins, at the level of both strains and functional genes, suggesting fine-scale resource partitioning in the human gut. No predictable responses to non-prebiotic micronutrients were found. Surprisingly, we did not observe decreases in day-to-day variability of the microbiota compared to a complex, varying diet, and instead found evidence of diet-induced stress and an associated loss of biodiversity. Our data offer insights into the effect of a low complexity diet on the gut microbiome, and suggest that effective personalized dietary interventions will rely on functional, strain-level characterization of a patient’s microbiota.

## Introduction

A quantitative understanding of the forces that shape the composition of the microbiota *in vivo* is a prerequisite to rationally engineering or manipulating it toward a favorable clinical outcome. Diet has been shown to play a pivotal role in shaping the microbiota from an early age. For example, a study of the fecal microbiota of children from a rural African village in Burkina Faso found significant enrichment in Bacteroidetes and depletion in Firmicutes compared to Western counterparts, which is associated with a diet enriched in complex polysaccharides ^1^. In the same vein, a dietary exchange experiment between a rural African cohort and an urban, African-American cohort, led to concomitant changes in their microbiota and disease-associated molecular biomarkers for the duration of the exchange ^2^. This indicates that commensal microbiota may reflect long-term dietary trends but remain labile in the face of dietary perturbations. Indeed, it was also shown that short-term, large-scale changes in diet can result in rapid and reproducible effects on subjects’ gut microbiome composition ^3^. Observations such as these highlight the interplay between a subject’s diet and their microbiota, and suggest that manipulating diet to engineer the gut microbiome may be a promising clinical intervention, together with rationally designed probiotics.

The interaction between short-term diet and the host microbiome is also an important and understudied confounder in many microbiome disease association studies. For example, both long-term dietary differences ^4^ and alterations of the microbiota ^5^ have independently been shown to be associated with differential rates of colon cancer; in the absence of a defined mechanism, it is unclear whether disease-associated microbial composition results from covariance with diet, which itself is causative, or whether the two factors are orthogonal and causally additive. A detailed and quantitative understanding of the impact of diet on the microbiome is therefore needed if the clinical vision of manipulating a patient’s diet to engineer their gut microbiome’s composition and/or metabolic output is to be achieved. However, identifying microbiome-diet interactions is complicated by a number of factors. Matching diets across patients is difficult, even if fixed meal plans are used, because participants may eat different quantities of individual components, altering the ratio of nutrients. Moreover, complex diets make it difficult to identify the role of individual micronutrients.

Previous studies have shown that microbiome composition responds to perturbations in diet ^3,6,7^ Indeed, some studies have shown that prebiotics affect the composition of the microbiota in predictable manner: for example, it has been shown that supplementation with inulin results increased abundance of *Bifidobacterium* and *Faecalibacterium* in a cohort of obese women ^8^, as well as *Lactobacillus* and *Bifidobacterium* in rats ^9^. A more recent double-blind, randomized, cross-over intervention study identified inulin-specific responses from *Anaerostipes, Bilophila* and *Bifidobacterium*^7^. However, perturbation studies in humans typically involve broad changes across complex mixtures of micronutrients ^3^, measure the response to a particular spike-in against a variable diet background ^8^, or perform relative abundance calculations from culture-based or qPCR methods ^10,11^. Resource partitioning in environmental bacteria has been shown to occur at exquisitely fine scales ^12^, yet little is known about resource partitioning in the human gut. If most species are generalists with respect to substrate use, then even a targeted addition can induce changes in many species, each competing for the same substrate. If most species are specialists, then we expect a strong response in a select few taxa.

Studies aimed at understanding how the microbiome will respond to a targeted change in a specific micronutrient remain logistically daunting. To overcome these challenges, we conducted a highly controlled feeding study and dietary perturbation experiment in a healthy human cohort, in which participants were placed on a standardized liquid nutritional meal-replacement for six days. Against this controlled dietary background, we investigated the effect of individual micronutrient spike-ins, including several prebiotic supplements, to identify the microbial responders and assess the extent to which prebiotics result in predictable compositional changes in the human microbiome.

## Results

### Study Design

To examine the response of the healthy human microbiome to targeted addition of micronutrients, we enrolled 60 healthy participants and randomized each to one of seven dietary spike-in arms: pectin, inulin, cellulose, fish oil (unsaturated fat), coconut oil (saturated fat), protein powder, and control (no spike-in). These spike-ins were chosen to cover the dominant nutrient categories typically present in a human diet, namely soluble fiber (inulin and pectin), insoluble fiber (cellulose), unsaturated fat (fish oil), saturated fat (coconut oil), and protein. Basic demographic data and the results of the randomization process are available in Table 1. In order to assess day-to-day variability within and between participants under their habitual, variable diet, we first collected two consecutive baseline stool samples, after which participants underwent a bowel cleanse using an over-the-counter osmotic laxative to remove remnants of their previous diet from their colon. The following day, they began the standardized diet period: for the first three days, participants consumed only a standardized liquid nutritional meal-replacement (Ensure Original, Abbot Nutrition) that is routinely used in both inpatient and outpatient care, and water. This was followed by three further days of liquid nutritional meal-replacement and water plus their assigned spike-in (Fig. 1a). The duration of three days for the initial equilibration period was chosen based on previous observations of autocorrelation dynamics in long-term time series of the gut microbiota in healthy humans, which found that most autocorrelations decay within 3 days ^13^. Participants were instructed to consume liquid nutritional meal-replacement *ad libitum* to allow for differential caloric requirements without compromising the relative composition of the standardized diet, and requested to submit stool samples preserved in nucleic acid stabilization buffer upon natural passage, with a maximum of one sample per day. Analysis of the average daily caloric intakes on days 3 and 6 did not show significant differences in caloric intake in a given arm. A post-intervention sample was taken on the first day of the resumption of routine, normal diet, as well as a follow-up sample one week later, to assess the persistence of any diet-induced compositional changes on the microbiota. All samples were processed within three days of receipt with only a single freeze-thaw cycle between sample processing and DNA extraction, library preparation and sequencing. Precise details of the standardized diet can be found in the Methods section.

**Figure 1.**
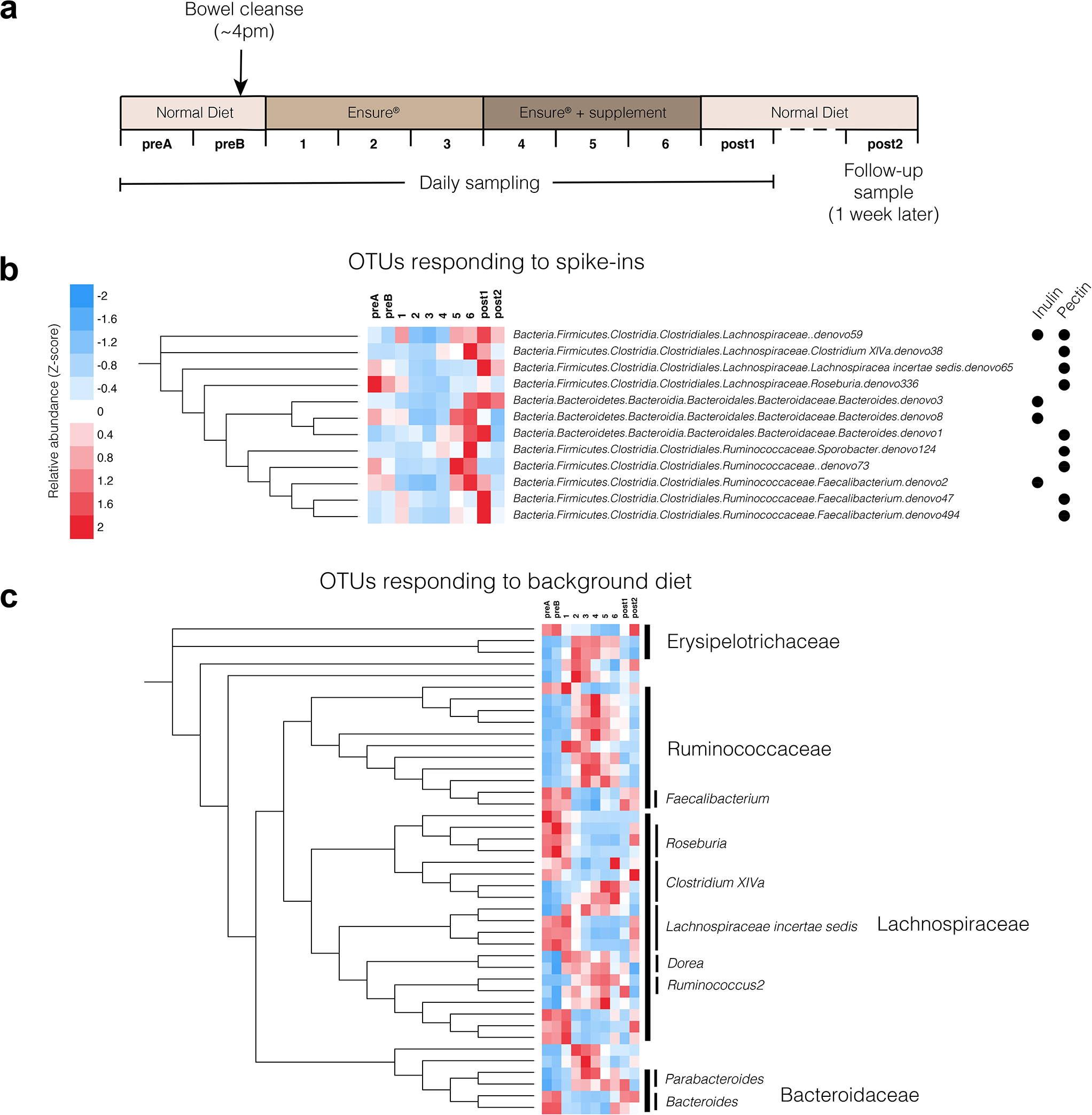
(a) Schematic outlining the dietary and sampling regimen for the study. (b) OTUs that showed statistically significant (DESeq2, FDR < 0.1) differential abundance on day 6 compared to day 3 in response to particular spike-ins. Mean relative abundances are depicted as Z-scores, computed across all participants. (c) OTUs that showed statistically significant (Wilcoxon Rank-Sum test, FDR < 0.1; *N*=39) differential abundance on day 3 compared to baseline. Complete RDP taxonomies can be found in Fig. S1.

**Table 1.** Demographic and baseline data for each spike-in arm.

| Arm | Number enrolled | Age (mean, range) | Number of bowel movements (baseline, day 6) | Average number of Ensure calories consumed (day 3, day 6) | BMI (mean, range) | Gender distribution (females, males) |
| --- | --- | --- | --- | --- | --- | --- |
| Inulin | 11 | 27.4 (23-30) | 1.0, 1.4 | 2156, 1975 | 23.6 (19-30) | 5, 6 |
| Pectin | 11 | 24.8 (23-28) | 1.3, 1.4 | 1728, 1920 | 23.5 (22-28) | 5, 6 |
| Protein | 5 | 25.2 (22-26) | 1.3, 1.2 | 1815, 1870 | 21.4 (18-25) | 2, 3 |
| Saturated fat | 7 | 24.4 (23-31) | 1.7, 1.6 | 2127, 2420 | 24.2 (19-27) | 2, 5 |
| Unsaturated fat | 6 | 23.3 (23-33) | 2.0, 1.2 | 1980, 2640 | 23.7 (20-29) | 2, 4 |
| Cellulose | 10 | 29.0 (27-31) | 1.3, 1.5 | 1886, 1860 | 24.1 (20-30) | 5, 5 |
| Control | 10 | 25.4 (22-32) | 0.8, 1.3 | 1760, 2310 | 22.4 (20-24) | 4, 6 |

### Response to nutrient addition

Targeted nutrient additions induced a targeted microbial response, suggesting a high-level of resource partitioning within the human gut. On average, a typical participant had 9 OTUs that exhibited a greater than tenfold cumulative gain in relative abundance over the spike-in period (including sample ‘post1’), and 14 OTUs with a fivefold gain. Some, but not all, nutrients yielded specific responses consistent across multiple individuals. To do this, we used differential abundance tests between day 3 and day 6 in each arm using DESeq2 ^14^ using raw 16S rRNA counts data (FDR < 0.1). DESeq2 has been shown to have high sensitivity on small sample numbers (*N*<20) ^15^. The only spike-ins with predictable and statistically significant responses across participants were pectin and inulin (Fig. 1b). This underscores their identity as prebiotics capable of altering microbial composition, as the other nutrients considered are typically absorbed in the small intestine and therefore not as likely to directly affect the composition of the colonic and stool microbiota. OTUs that responded to pectin or inulin generally reached their peak between day 5 and the first day of resuming a normal diet (post1). This is consistent with typical transit times of food in the gut, which vary by person but have been previously shown to involve a period of approximately 24–48 hours by using food dyes ^3^. Responders to the prebiotic spike-ins were limited to the Clostridial *Lachnospiraceae* and *Ruminococcaceae* families, and the *Bacteroides* genus. While taxonomies were assigned using the RDP classifier ^16^ and therefore did not include species-level annotation, we performed BLAST searches of the centroid 16S rRNA sequences for OTUs of interest against the NR database on NCBI. Notably, members of the same species appeared to respond to different spike-ins (Fig. 1b): in particular, the 16S rRNA sequences of the *Faecalibacterium* OTU responders, one of which responded to inulin and two to pectin, all mapped with 100% identity to *Faecalibacterium prausnitzii* genomes, suggesting that different commensal strains of the same species have different carbohydrate-active enzyme specificities. This is consistent with previous studies that reported *F. prausnitzii* blooms *in vivo* in participants given inulin as a supplement ^17^, and also that commensal *F. prausnitzii* isolates can be cultured *in vitro* on pectin ^18^.

Importantly, certain OTU responders exhibited highly predictable and reproducible behavior across people. In particular, a *Bacteroides* OTU that mapped with 100% identity to *B. uniformis* genomes on the NR database bloomed strongly in inulin-consuming participants, reaching relative abundances between four- and sixteen-fold greater than that on day 3 in the same participant after consuming the spike-in (Fig 2a). In contrast, a number of OTUs appeared to respond to specific spike-ins in some participants but not others. These were undetectable using a DESeq2-based differential abundance test across all participants in the given arm, but were identifiable through an analysis of the individual trajectories between days 3 and post1. In particular, the cellulose arm was significantly enriched with these person-specific responses. These responses can take two forms: in the first kind, the bacterial responder was present in most or all participants, but only bloomed in a subset of them, while in the second kind, the responder was not present in all participants but when it was, bloomed reproducibly. The most notable example of the first kind was an OTU which mapped with 100% identity to *Bacteroides cellulosilyticus*, a known cellulose degrader and commensal inhabitant of the human gut ^19^, which exhibited an extremely strong response in a single person, despite being present in all other participants within that arm (Fig. 2b). In contrast, Archaeal methanogens were only present in a subset of people, but in those participants bloomed in response to cellulose (Fig. S2). This finding is consistent with previous observations that only a subset of humans are known to harbor commensal Methanobacteria, which utilize hydrogen produced by certain bacterial cellulose degraders for methanogenesis ^20,21^. It is also relevant in the context of reducing the net microbial production of hydrogen (as opposed to methane), which is one of the bases for a low FODMAP diet in the context of treating Irritable Bowel Syndrome ^22^.

**Figure 2.**
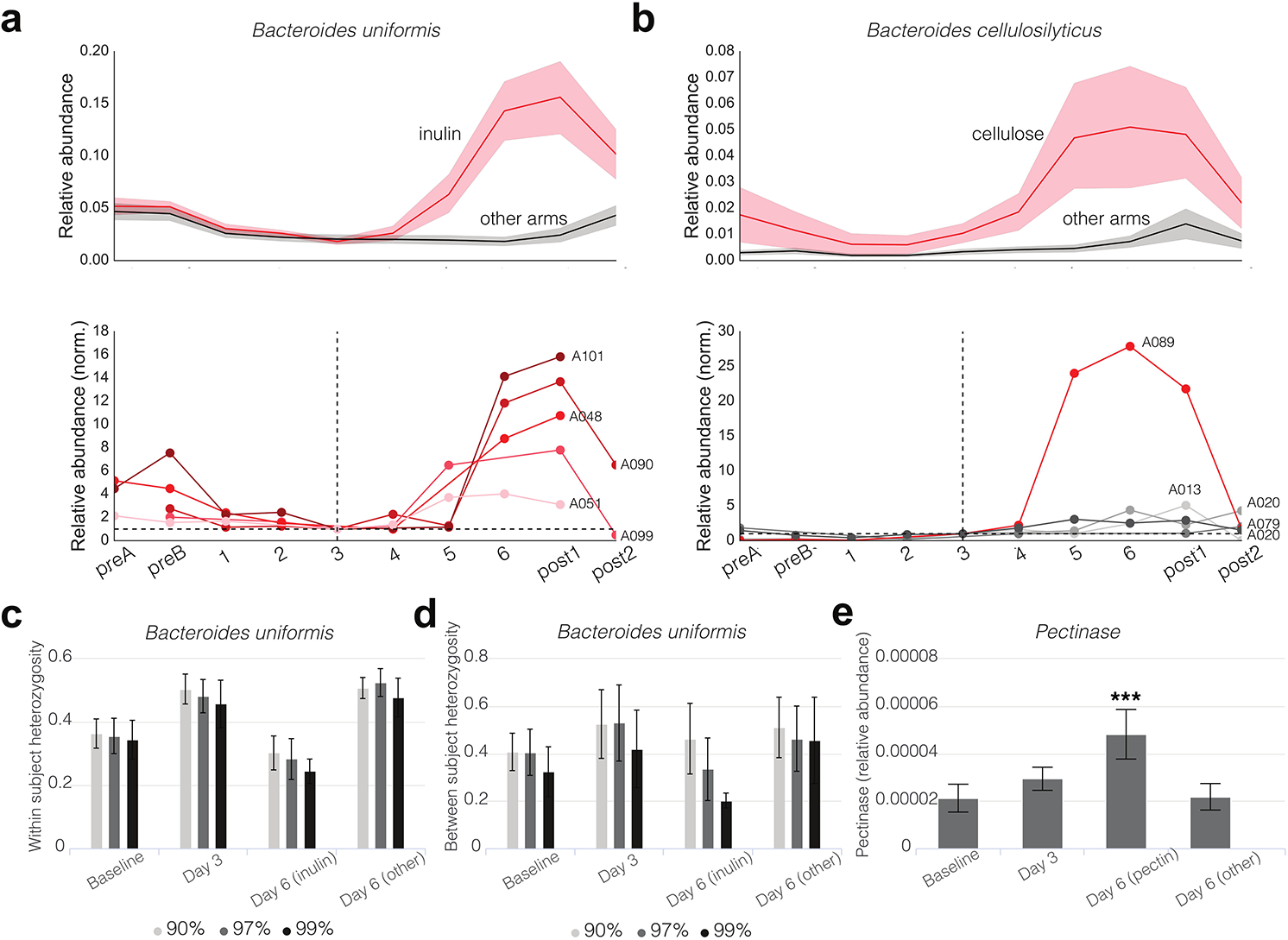
(a) *Bacteroides uniformis* blooms in response to inulin supplementation across all participants. The top plot shows mean and standard deviations of relative abundances across participants, and the bottom shows individual participant timeseries, normalized to relative abundance on day 3 for comparison purposes. (b) Same visualization for *Bacteroides cellulosilyticus.* (c) Average within-participant SNP heterozygosity in AMPHORA genes of *B. uniformis*. (d) Average between-participant SNP heterozygosity in AMPHORA genes of *B. uniformis*. (e) Relative abundance of pectinase enzymes in the metagenome. Statistical significance of the difference in abundance of pectinases on day 6 in the pectin arm was computed by a Wilcoxon Rank-Sum test compared to the abundance of pectinases on day 6 in the other arms (p<10^−5^).

### Strain-level dynamics

It is now known that polysaccharide specificity occurs at the species level ^23^. For example, species within the *Bacteroides* exhibit remarkable diversity in polysaccharide utilization ^24,25^. However, it is unclear to what extent individual strains exhibit differential polysaccharide utilization phenotypes. To investigate this point, we computed the mean heterozygosity of Single Nucleotide Polymorphisms (SNPs) within different species as a proxy for strain-level diversity (cf. Methods). The only bacterium that exhibited statistically significant differences in heterozygosity on day 6 compared to day 3 was *B. uniformis,* which had substantially lower heterozygosity in participants in the inulin arm on day 6 (Fig. 2c). This signal was retained (and indeed, stronger) as the identity cut-off for read mapping was increased up to 99%. This lower heterozygosity combined with increased relative abundance of the species in participants receiving inulin supplementation suggested that a specific strain of *B. uniformis* was being enriched in response to inulin.

Upon performing the same heterozygosity computation for *B. uniformis,* but this time between participants (i.e. the probability that overlapping read pairs, one from each person, exhibited a different allele), we observed a similar decrease in heterozygosity when a 97% and 99% identity cut-off was used, suggesting the same strain was being enriched across different participants (Fig. 2d). These data indicate that different strains of *B. uniformis* have different inulin-utilization phenotypes.

### Functional gene content

Since *B. uniformis* has a known inulin-specific polysaccharide utilization locus (PUL) in its genome ^24^, we reasoned that the enrichment of different OTUs or strains could be attributed to the presence of specific genomic carbohydrate active enzymes, and so mapped the metagenomic reads against HMM profiles for inulinases and pectinases in dbCAN ^26^. Consistent with this hypothesis, we observed an enrichment of pectinases in the metagenomes of participants in the pectin arm on day 6 (Fig. 2e). Surprisingly, however, we did not observe any such differences for inulinases in the inulin arm, perhaps due to poor annotation and characterization of these glycoside hydrolases and PULs in the human gut metagenome.

### Response to background diet

Having shown that the response to a targeted dietary perturbation was both specific to the spike-in and in some cases generalizable across participants, we next turned our attention to the large-scale dietary perturbation built in to our study design: the switch to a standardized diet. We first looked for taxa that reproducibly responded to the diet change across participants by identifying OTUs that were differentially abundant on day 3 compared to baseline using a Wilcoxon Rank-Sum Test (FDR < 0.1). OTUs that responded to the dietary perturbation generally fell into two clearly identifiable groups which followed two opposing dynamic trajectories (Fig. 1c and Fig. S1): one group of OTUs increased in abundance on nutritional meal-replacement compared to baseline, only to become depleted once a normal diet was resumed, and a second group of OTUs became depleted on nutritional meal-replacement relative to baseline, only to return upon resumption of normal diet. The majority of OTU responders in both groups belonged to the Clostridial bacterial families *Ruminococcaceae* and *Lachnospiraceae,* with some exceptions (most notably *Erysipelotrichaeceae, Deltaproteobacteria, Parabacteroides* and *Bacteroides*). Interestingly, there was no apparent high-level phylogenetic association with either group: indeed, OTUs assigned to the same bacterial genus by RDP sometimes exhibited opposing dynamics (e.g. OTUs in *Clostridium XIVa*).

Since the specific liquid nutritional meal-replacement used in this study is frequently used by physicians, dieticians and healthy individuals in a clinical context, our dataset offered a unique perspective on its effect on the microbiota of otherwise healthy humans. Aggregating participants from all spike-in arms, we counted the number of OTUs present and absent before and after the study, and found that participants lost an average of 32.3% of their total number of baseline 16S OTUs during the period of fixed diet, only 16.1% of which were regained after a week of resuming a normal diet. It is possible that the bowel cleanse was responsible for this loss, so to test the extent to which the cleanse might be able to explain the loss of diversity, we reasoned that organisms that were still present on day 2 of the liquid nutritional meal-replacement were unlikely to have been lost as direct result of the cleanse alone. Of the OTUs lost by the end of the prescribed diet, approximately half (17.6%) were lost by day 2, and the remainder was lost after day 2. Of course, even for the latter organisms, we cannot eliminate the possibility that the cleanse did not act in concert with the diet as a driving force for their disappearance. Nonetheless, these data suggest a loss of approximately 16% in the measurable number of OTUs. We then repeated the SNP-level heterozygosity analysis described above for all species in our reference genome set, and obtained mean heterozygosity values across all AMPHORA genes in all species for a given participant (Fig. 3a). We found that overall, the mean heterozygosity of SNPs in the entire microbiota increased significantly from baseline to day 3, a change that was maintained (although slightly diminished) by day 6. The effect for *Faecalibacterium,* an important commensal organism that has been negatively associated with Crohn’s disease ^27^, was particularly strong (Fig. 3b). This relative increase in the strain-level diversity of the microbiota suggested a flattening of the fitness landscape between strains. This observation is reminiscent of previous work in *E. coli* that showed certain environmental stresses can alleviate the effect of deleterious mutations and therefore reduce fitness differences between strains ^28^.

**Figure 3.**
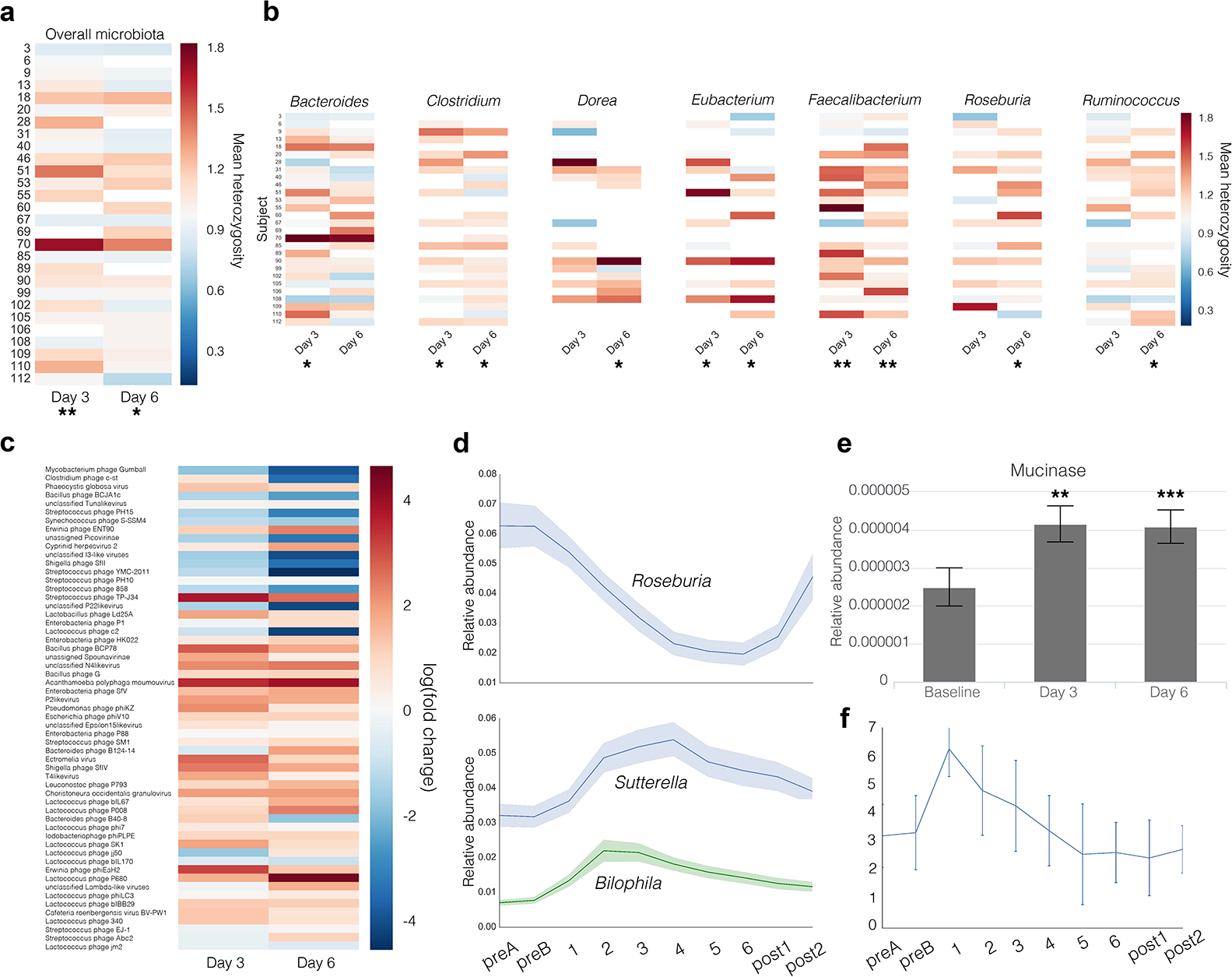
(a) Mean AMPHORA gene SNP heterozygosities within each participant on days 3 and 6, averaged across all bacterial species and normalized to baseline levels. (b) Mean AMPHORA gene SNP heterozygosities averaged across the most abundant genera in the gut. (c) Log-transformed mean relative abundance of various bacteriophages in all participants, normalized to baseline levels. (d) Mean and standard error timeseries of the relative abundance of *Roseburia, Bilophila* and *Sutterella.* (e) Relative abundance of mucinase genes at baseline, day 3 and day 6. (f) Bristol Stool Scale of all participants throughout the study.

We hypothesized that if the dietary change led to an increase in stress for the bacterial hosts, then we might expect to see an increase in active phage. The relative abundance of bacteriophage DNA in metagenomic sequences on days 3 and 6 relative to baseline (Fig. 3c) suggested that many lysogenic phages were entering the lytic phase of their life cycles in response to the dietary perturbation.

The most striking and reproducible effect of the nutritional meal-replacement diet on microbiome composition was a decrease in the relative abundance of members of the *Roseburia* genus (Fig. 3d), an important and abundant butyrate-producing, mucosa-colonizing commensal that has been shown to be negatively associated with Crohn’s disease and ulcerative colitis, types of inflammatory bowel disease (IBD) ^29,30^. Concomitantly, a transient increase in Proteobacteria was observed: specifically, the genera *Bilophila* and *Sutterella* were found to transiently increase in relative abundance during the diet, peaking on days 2 and 4, respectively, before slowly returning to their baseline levels. While the physiological effect of these changes on the host is difficult to assess from genomic data alone, it is worth noting that both of these Proteobacterial genera have potential associations with disease: *Bilophila* has been associated with the production of sulfide, which leads to degradation of host mucin and induces colitis in mouse models ^31,32^, while *Sutterella* has been associated with insulin resistance in obese patients, and autism with gastrointestinal disturbance in children ^33,34^.

At the functional level, we observed significant increases in the relative abundance of mucinase genes in participants’ metagenome in on days 3 and 6 relative to baseline (Fig. 3e), consistent with a diet significantly impoverished in dietary fibers and complex polysaccharides, in which the colonic microbiota rely on host mucins as a carbon source ^35^. As in the mouse models that first documented this observation, the effect was not rescued by any of the individual prebiotic spike-ins ^35^. The increase in mucinases may also be linked to the depletion of *Roseburia* noted earlier: as a mucosal inhabitant, it is likely that destruction of the host mucin disrupted its native ecological niche. Importantly, loss of *Roseburia* was partially reversed one week after resuming normal diet (Fig. 3d), suggesting that the resumption of a more complex diet for a period of one week was able to partially restore the mucosal layer and its *Roseburia* inhabitants.

To quantify the extent of change induced by diet as a function of time, we computed a weighted Unifrac distance between participant pairs before and after the prescribed diet. We found that participants resembled each other more closely on day 6 than they did at baseline, but only marginally so. However, the effect was stronger in the prebiotic spike-in arms (PERMANOVA, *p*=0.04), suggesting that prebiotics led to a more predictable and reproducible response across participants compared to the standardized nutritional meal-replacement diet (Fig. 4a). We reasoned that this could be explained by the chemical nature of the diet: in the case of prebiotics, only a small set of organisms appeared to respond, but their responses were consistent across people (Fig. 1b and 2a); in contrast, though the chemical composition of the standardized liquid diet was identical in all participants in the study, it was dominated by processed sugars, which are primarily absorbed in the small intestine, whose remainders act as ubiquitous substrates that would be chemically accessible to a larger fraction of the microbiota, thereby increasing stochasticity. We hypothesize that large inputs of non-specific substrates affect each participant’s community differently and as a function of their initial composition.

**Figure 4.**
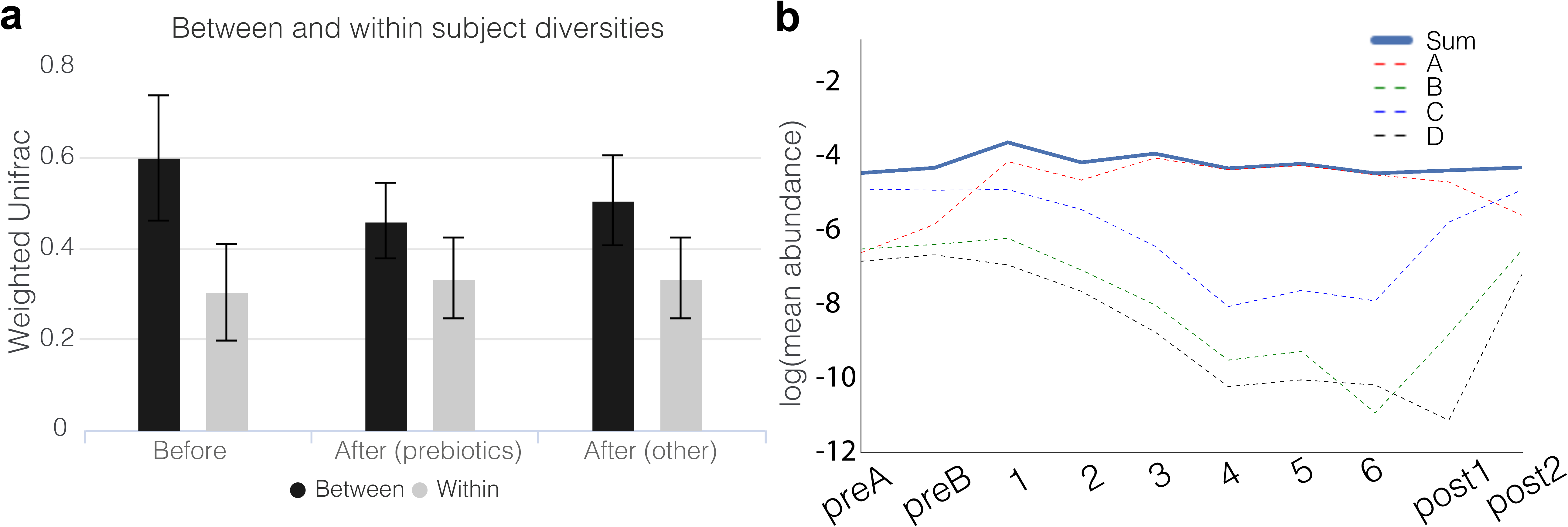
(a) Weighted Unifrac values between and within participants at baseline (before) and day 6 (after). Error bars represent standard deviations. (b) Log-transformed mean relative abundance of the four distinct OTUs given a taxonomic assignment of *Lachnospiracea incertae sedis* at the genus level and that were found to be differentially abundant on day 3 compared to baseline. Their combined relative abundance is shown in a solid line. For clarity, the *denovo* OTU IDs are labeled as follows: *denovo6* - A; *denovo84* - B; *denovo65* - C; *denovo155* - D.

Previous studies showed extensive day-to-day variation in the gut microbiome ^36^, hypothesizing that a large fraction of this variance may be due to diet ^13^. We asked whether keeping diet standardized would reduce daily variation. We computed Weighted Unifrac distances between adjacent days within a person during a period of variable diet (samples ‘preA’ and ‘preB’) and compared this with Weighted Unifrac distances within the same participant on adjacent days in which the diet was identical (days 5 and 6). To our surprise, we found that there was no discernible increase in day-to-day similarity on a completely identical diet, irrespective of whether participants were given prebiotic spike-ins (Fig. 4a). These data suggest that short-term dietary fluctuations on daily timescales contribute in an only very limited manner to the day-to-day variability observed in a participant’s fecal microbiota, and that other variables, including fluctuations in host lifestyle, physiology, immunology, and perhaps even experimental noise are more important factors.

We found that a participant’s sample could reliably be classified as pre- or post-nutritional meal-replacement. Two Random Forest Classifiers (RFCs) were constructed to classify samples as either baseline or post1 (the first day after Ensure), and baseline or post2 (one week after resuming a normal diet), respectively. In the first classifier, we obtained an AUC of 0.93 (*p*<10^−7^), while in the second, we obtained an AUC of 0.90 (*p*<10^−5^) (Fig. S3). These data indicate that there remain effects of the fixed diet on the overall microbiota even one week after resuming a normal diet (Table S2). The top 10 features of the baseline/post1 RFC included three of the four OTUs assigned to *Lachnospiraceae incertae sedis* that appeared in Fig. 1c. Digging deeper, we found that these OTUs were indeed depleted on nutritional meal-replacement, while the remaining OTU replaces them, resulting in a constant combined relative abundance through time (Fig. 4b). These dynamics suggest a switch in occupancy of relative organisms in a particular niche in response to the dietary perturbation.

## Discussion

When designing our study, we hypothesized that by introducing a constant diet background, we would improve our ability to detect microbial responders to particular prebiotic spike-ins. Indeed, we were able to detect reproducible microbial responses to both inulin and pectin across participants in relatively small numbers for each arm (<10) after filtering for early withdrawals or insufficient sampling. These responses were consistent across participants, including at the level of strains in the case of *B. uniformis*, indicating a strong predictability of the effects of prebiotic supplementation on the microbiota. Interestingly, we did not observe significant blooms of *Bifidobacterium* in response to inulin, which have previously been documented in the literature ^10,11,17,37^. While we cannot conclusively explain this discrepancy, it is worth noting that many such studies are culture-based, emphasizing the need for further study and verification of these phenomena using modern sequencing technologies directly applied to stool. It is also worth noting that the three *Bifidobacterium* OTUs present in the dataset were only present in a subset of the participants in the inulin arm, which necessarily reduced our statistical power to detect responses thereof. In the case of at least one such non-culture-based analysis of the Bifidogenic nature of inulin, subjects were specifically chosen to contain appreciable amounts of Bifidobacteria ^17^. This highlights the difficulty in observing statistically significant, reproducible responses of the microbiota *in vivo* in a cohort containing randomly chosen participants who may or may not contain the bacteria in question.

Moreover, in contrast to the predictability of inulin- and pectin-specific responses, certain responses were not detectable through traditional statistical means, because they only occurred in a subset of participants. The most noteworthy example was *B. cellulosilyticus*, which appeared to bloom strongly in response to cellulose in only one person. Archaeal methanogens also bloomed in responses to cellulose, but only in participants in which they were present. It is possible that these two cases are one and the same, i.e. that the *B. cellulosilyticus* OTU contains multiple strains with different functional properties, and the strain capable of degrading cellulose was present only in that one participant. Alternatively, it is possible that these person-specific blooms are manifestations of complex microbial networks on which this species depends to exhibit this phenotype. More broadly, these data offer a tantalizing glimpse into the potential for personalized nutritional interventions that account for strain-level composition of a patient’s microbiota.

We also hypothesized that standardizing diet across participants would reduce the quantitative contribution of short-term dietary fluctuations on the microbiota, a concern in the design of clinical studies aiming to investigate the association between microbial composition or fluctuations and disease. However, by placing all participants on a standardized, liquid diet consisting of a nutritional meal-replacement, we found that the diet introduced an axis of variation in the microbiota that did not appear to reach full stationarity after 6 days, and was further complicated by the difficulty of adhering to a liquid diet. From a practical standpoint, these data indicate that standardizing diet in a clinical cohort in this manner will have limited returns in reducing noise introduced by short-term dietary fluctuations, and may even introduce additional confounders. Averaging over additional participants or multiple timepoints may be more suitable means for reducing the impact of diet on other studies.

Moreover, the prescribed diet appeared to result in diet-induced stress on the participants’ microbiota, with a flattening of the overall fitness landscape at the level of strains, an increase in the relative abundance of a large number of phages, and increases in mucinase abundance in the metagenome, indicative of increased consumption of host mucin as a carbon source and degradation of the mucosal layer by the microbiota. While the diet-induced stress is potentially confounded with stress induced from the bowel cleanse, the reported changes persisted from days 3 to 6, well after bowel movements returned to baseline consistency (Fig. 3f), indicating that the prescribed diet was a likely causative factor underlying these observations. While we did not experimentally measure changes to the mucosa, it is worth noting that thinning of the defensive mucosal layer is associated with a number of intestinal diseases, including colon cancer and IBD ^38–40^. It is likely that the chemical contents of the nutritional meal-replacement we used, which consist to a large extent of processed sugars easily converted to glucose, can explain these changes. The ubiquity of this substrate likely shrinks the phenotypic differences between strains, and favors fast growing organisms over the more slowly dividing host commensals, as suggested by the Proteobacterial responders (Fig. 3d). Taken together, these data suggest that clinical use of dietary supplementation or meal replacement strategies would benefit from more thoughtful formulations designed with the microbiota in mind, by including a higher content of dietary fibers and complex polysaccharides selected for their prebiotic qualities, in order to minimize potentially detrimental effects to the host.

## Methods

### Experimental model and subject details

60 healthy human volunteers, consisting of 25 females and 35 males, were enrolled into the study under the supervision of the MIT Committee on the Use of Humans as Experimental Subjects (COUHES), who approved the study under protocol number 1504007066A002. All participants provided written, informed consent, and the study was conducted in accordance with the relevant guidelines and regulations. To be included, participants had to be between 18 and 70 years of age and have a BMI between 18 and 30. Exclusion criteria were food allergies or dietary intolerances of any kind, a history of irritable bowel syndrome, inflammatory bowel disease, Type-2 diabetes, kidney disease or intestinal obstruction, allergy to polyethylene glycol, antibiotics treatment in the 6 months leading up to the study, untreated *in situ* colorectal cancer, or currently pregnant, planning to get pregnant in the next 60 days, or breast-feeding. Baseline cohort data and the results of the randomization process are available in Table 1.

### Method details

#### Study logistics

Upon being consented into the study, participants were first randomized to a spike-in arm and were then provided with a kit consisting of materials for a bowel cleanse, sufficient stool collection hats and labeled Para-Pak vials (Meridian Biosciences, Inc.) with 5ml RNALater (Ambion, Inc.) for daily sampling, in addition to instructions for appropriate sampling strategy, and three daily portions of their allocated spike-in. They were also provided with several locations in which stool samples could be dropped off and additional liquid meal-replacement could be picked up, and were instructed to collect samples whenever they naturally passed stool, with at most one sample per day. Samples were collected daily by study coordinators at all drop-off locations, and transported to a Biosafety Level 2 laboratory at MIT, where they were processed daily in batches, and no later than three days after passage. In addition, participants completed a daily questionnaire to record Bristol Stool Scale and bowel movement frequency, report any notable changes in health that did not relate to the study, including consuming antibiotics and medication, and inform study coordinators of lapses in adherence to the standardized diet, as well as document the amount of liquid meal-replacement consumed that day. Samples obtained after any reported lapses were excluded from the analysis.

#### Dietary regimen

All participants included in the analysis followed the dietary regimen outlined in Fig. 1a. For two days prior to beginning the study, participants were instructed to continue their habitual diet and provide two baseline stool samples. After obtaining the second baseline stool sample, participants underwent a bowel cleanse in the evening by consuming 150g of over-the-counter osmotic laxative PEG 3350 (MiraLAX, Bayer) and 64oz bottles of electrolyte solution (Gatorade, PepsiCo) and were instructed to consume only clear liquids during the cleanse. Upon waking (day 1), participants began the standardized diet consisting entirely of liquid, nutritional meal-replacement (Ensure Original, Abbott Nutrition) and water for a period of six days. They were allowed to consume both *ad libitum* to avoid unnecessary weight loss and allow for heterogeneity in basal metabolic rates and daily caloric requirements. The liquid, nutritional meal-replacement was available in several flavors: vanilla, chocolate, strawberry, butter pecan, and coffee latte. Dark chocolate was excluded due to an increased amount of dietary fiber compared to the other flavors, which shared the same composition. Participants were allowed to choose freely between flavors, but were requested to maintain a consistent daily combination if they intended to consume several flavors. Nutritional composition of a single serving (one 8oz bottle) of the liquid, nutritional meal replacement reported by Abbott Nutrition is as follows: 220 calories, consisting of 6g total fat (1g saturated fat, 0g trans fat, 2g polyunsaturated fat, 3g monounsaturated fat), <5mg Cholesterol, 200mg Sodium, 370mg Potassium, 32g total carbohydrate (<1g dietary fiber, 15g sugars, derived mostly from corn maltodextrin and glucose), and 9g protein. This consists of a diet that is significantly impoverished in dietary fiber, which we reasoned would improve our ability to detect changes in response to prebiotic spike-ins.

Participants were also provided with three daily doses of their allocated spike-in, to be consumed on days 4, 5 and 6. Spike-ins were given in pre-weighed powder form for inulin, pectin, cellulose and protein powder mixture, while fish oil was provided in pill form, and coconut oil was provided raw in pre-weighed tubes. Participants were instructed to consume their spike-ins throughout the day by mixing them either with a serving of liquid meal-replacement or water, and to record any lapses in either adherence to the standardized diet or to the consumption of their allocated spike-in on the daily questionnaires provided. The allocated daily doses of each spike-in were as follows: 10g/day of inulin (CAS number 9005-80-5, Alfa Aesar); 35g/day of pectin (CAS number 9000-69-5, MP Biomedicals, LLC); 20g/day of cellulose (NutriCology); 40g/day coconut oil (Kirkland Signature Organic Coconut Oil); 6 pills of fish oil each containing 1050mg of omega-3, -5, -6, -7, -9 and -11 unsaturated fatty acids (Kirkland Signature Wild Alaskan Fish Oil); 51g of protein powder, consisting of 25.5g of whey protein (Optimum Nutrition Gold Standard 100% Whey Protein Powder) and 25.5g of casein protein (Optimum Nutrition Gold Standard 100% Casein Protein Powder); no spike-in for control. Spike-in doses were set at 150% of the Recommended Daily Allowance (RDA) for that particular nutrient, where available. In the absence of a RDA (e.g. inulin, pectin, and cellulose), doses were set at the highest dose that the investigators determined could reasonably ingested by a participant over the course of a day without experiencing discomfort.

Participants were instructed to consume the standardized diet plus allocated spike-in until the end of day 6. Two follow-up samples were also obtained, one on the day of resuming a normal, variable diet (post1) and another a week later (post2).

#### Stool sample processing and DNA sequencing

Participants collected approximately 1g of stool and dissolved it in 5ml RNALater (Ambion, Inc.) pre-aliquoted into Para-Pak vials (Meridian Biosciences, Inc.). In order to maximize our ability to obtain clean genomics signals and avoid experimental and sample processing confounders, great efforts were taken to ensure that all samples were treated in the same manner. Samples were picked up from daily drop-off locations by study coordinators daily, and washed with PBS the same day or no longer than 3 days upon passage before being frozen at −80°C. 29 of the 60 subjects returned a complete set of daily stool samples for the entirety of the requested timeseries. Samples were aggregated through time and submitted together for DNA extraction, library preparation and DNA sequencing at the Genomics Platform at Broad Institute of MIT and Harvard. 16S rRNA regions were amplified from all samples using a universal V4 primer. Baseline, day 3 and day 6 samples were also sequenced using shotgun metagenomics sequencing. 16S rRNA amplicon sequencing was performed on an Illumina MiSeq using v2 chemistry, and metagenomics sequencing was performed on an Illumina HiSeq2500 High Output flowcell using v4 chemistry.

### Quantification and statistical analysis

#### 16S rRNA sequencing data analysis

##### 1. Raw read processing and OTU calling

Raw paired-end 16S rRNA Illumina sequencing reads were merged with PEAR ^41^, resulting in a total of 64,743,565 raw merged reads. Reads were then demultiplexed, and quality trimmed with a cut-off of Q=25 using usearch8 ^42^, before being trimmed to a common length of 226 bases. Dereplicated reads were then clustered into OTUs to 97% identity using UPARSE ^43^. OTU centroids were assigned a taxonomy using the RDP classifier using an uncertainty cut-off of 0.5 ^16^. Since RDP does not return species annotation, we also performed BLAST searches of the centroid 16S rRNA sequences for OTUs of interest against the NR database on NCBI ^44^. A species annotation was given if the top 10 alignments matched a species’ genome with 100% identity. Samples with fewer than 5,000 reads (totaling 38 out of 501) were discarded from the analysis.

##### 2. Phylogenetic trees and distance calculations

A phylogenetic tree was constructed from all 16S OTU centroid sequences using FastTree ^45^. Sub-trees in Figure 1 were computed in the same manner but using only the OTU sequences of interest, and plotted using iTOL ^46^. Beta diversity calculations between samples (Fig. 4a) were computed from rarefied OTU tables with the Weighted Unifrac distance ^47^ using the scikit-bio Python package and the phylogenetic tree mentioned in the section above. Prior to computing beta diversity values, a rarefaction was performed on the OTU table by down-sampling each sample to 5,516 reads (the minimum read count in the dataset after filtering), to remove artifactual effects of differential read counts across samples. Statistical significance of the differences in between-subject beta diversities on day 6 in the prebiotic versus non-prebiotic arms was assessed with a PERMANOVA test (*N*=29, 10,000 permutations) using the scikit-bio Python package.

##### 3. Statistical tests of differential abundance

Statistical tests to identify OTU responders to spike-ins (Fig. 1b) were performed using DESeq2 ^14^. Participants who had both day 3 and day 6 samples were aggregated into their respective spike-in arms, and the test was performed between those two days. Due to the effects of attrition on participant numbers and since not all participants produced daily stool, day 5 samples were also included in the day 6 category in the absence of a day 6 sample. An FDR cut-off of 0.1 was used to identify responders to a spike-in, provided it did not also meet this threshold in the control arm. The number of participants that matched these criteria in each arm was N=9 pectin, N=5 inulin, N=4 cellulose, N=6 in the saturated fat, N=3 unsaturated fat, N=2 protein, and N=6 control.

OTU responders to the standardized, nutritional meal-replacement background (Fig. 1c and S1) were identified using a Wilcoxon rank-sum test between the latest baseline sample and day 3 (N=39), with a Benjamini/Hochberg FDR cut-off of 0.1.

#### Metagenomics sequencing data analysis

##### 1. Raw read processing

Human reads were removed from raw forward and reverse Illumina Whole Genome Shotgun (WGS) reads by aligning to the hg18 human genome from the UCSC Genome Browser ^48^ using BWA ^49^. To remove biases introduced by PCR replicates, remaining reads were then dereplicated using PRINSEQ ^50^, for all exact matches and 5’ duplicates (prinseq option: -derep 2). Unique reads were then quality-trimmed with trimmomatic to Q=20 ^51^, to a final total readcount of 1,695,900,645, or an average sequencing depth of 15,007,970 reads per sample, with 15 out of 113 samples having fewer than 1,000,000 reads.

##### 2. Carbohydrate active enzyme annotations

Functional annotations for reads were obtained from the CAZY database using dbCAN ^26^. Pre-processed metagenomics reads were converted to the 6 possible Open Reading Frames using using EMBOSS Transeq ^52^ and aligned to the HMM profiles using HMMER3.0 ^53^. The profiles searched were: GH32, GH91 and CBM38 for inulinases/fructanases, and CE8, CE13, PL1 and GT47 for pectinases. Only CE8 (shown in Fig. 2e) showed statistically significant differences as measured by Wilcoxon rank-sum tests on the relative abundances on days 3 and day 6. We also searched PL8 for hyaluronate lyases (EC 4.2.2.1), also known as ‘mucinases’ (as shown in Fig. 3). A 1e-3 E-value cutoff was used for alignments less than 80 amino acids long, and 1e-5 for longer alignments.

##### 3. Single nucleotide polymorphism (SNP) heterozygosity calculations

Heterozygosity values reported in Figures 2 and 3 were computed using AMPHORA ^54^ to identify thirty-one single-copy phylogenetic markers in a set of 649 non-redundant reference genmes from the Human Microbiome Project ^55^. Metagenomics reads were mapped from each sample to these genes using BWA-mem ^49^ with the “-a” flag and a conservative cut-off of 90% sequence identity, which in previous work has been shown to be well under the species boundary for AMPHORA genes (Smillie, *in press* 2017). These alignments were then filtered for SNPs by removing all monomorphic sites. Next, all samples with zero coverage at more than 25% of the identified SNP loci were removed, and finally alignment sites with atypical coverage, defined as being greater than 1.5 standard deviations away from the mean coverage, were also removed. Within-subject heterozygosity values were computed at each site by considering all read pairs within a sample overlap at that site, and computing the probability that any read pair had the same allele. From the Hardy-Weinberg equilibrium, 1 minus this value gives the heterozygosity. Between-subject heterozygosity values were computed in the same fashion, except that all overlapping read pairs containing a common SNP locus were enumerated using reads coming from pairs of subjects, i.e. one read per subject. Python code for performing these calculations from sorted BAM files can be found at the following Github link: https://github.com/thomasgurry/strains/blob/master/heterozygosity.py

##### 4. Bacteriophage annotations

Relative abundance of phages were estimated using Kraken and the full NCBI phage database ^56^. Filtered read counts were converted to relative abundances by dividing by the total number of reads.

#### Machine learning

Random Forest Classifiers (RFCs) were built using the scikit-learn Python package. Classifiers were built using subjects which contained one of each sample of the classification classes: for example, a subject had to contain one of ‘preA’ or ‘preB’ samples and a ‘post1’ sample to be included in the baseline/post1 RFC. This ensured that each class was balanced and that no biases could be introduced from inclusion of samples from a subject in only one of the two classes. The baseline/post1 and baseline/post2 RFCs had *N*=26 and *N*=23 in each class, respectively. Cross-validation was performed using a ‘leave a subject out’ approach, i.e. by successively removing both samples from a subject samples and using them as test sets. P-values for each classifier were computed using Fisher exact tests on the resulting confusion matrices.

## Acknowledgements

The authors wish to thank Dr. Ashley Vargas from the NIH for preliminary feedback on the study design, Dr. Vicki Mountain and Paige Swanson for logistical support, Dr. Jason Zhang for helpful discussions, and the Rasmussen family foundation for funding. This project was in part supported by award Number 132GM007753 from the National Institute of General Medical Sciences. The content is solely the responsibility of the authors and does not necessarily represent the official views of the National Institute of General Medical Sciences or the National Institutes of Health.

## Author Contributions Statement

T.G. designed the study, performed participant recruitment and sample processing, analyzed the data and wrote the manuscript. HST.M.C. designed the study, performed participant recruitment, and reviewed the manuscript. S.G. assisted in analyzing the data. L.T.T.N. assisted in study design and sample processing. S.M.K. assisted in data analysis. A.A. reviewed study design and acted as study physician. X.J. assisted in data analysis. Z.K. assisted in study design and writing the manuscript. E.J.A. designed the study, analyzed the data and wrote the manuscript. All authors reviewed the manuscript.

## Additional Information

### Data Availability Statement

The datasets generated during and/or analysed during the current study are available from the corresponding author on reasonable request.

